# Spinal cord pathology in a Dravet Syndrome mouse model

**DOI:** 10.1101/2023.09.22.558962

**Authors:** Juan Antinao Diaz, Ellie Chilcott, Anna Keegan, Stephanie Schorge, Simon N Waddington, Rajvinder Karda

**Author notes:** Corresponding Author: Rajvinder Karda, 86-96 Chenies Mews, London, WC1E 6HX, UK.

## Abstract

**Summary:** *Objectives:* Dravet syndrome is a severe epileptic encephalopathy that begins in early childhood. More than 80% of patients with Dravet syndrome exhibit a haploinsufficiency in *SCN1A*, which encodes the voltage-gated sodium ion channel Na_V_1.1. The epilepsy is believed be caused by specific deficit of *SCN1A* in inhibitory interneurons of the hippocampus. However, the aetiology of other symptoms including gait disturbances, ataxia, cardiac issues and dysautonomia is less clear.

*Methods:* In an *Scn1a* knock-out (*Scn1a*^-/-^) mouse model which recapitulates clinical phenotypes, we assessed Na_V_1.1 and neuroinflammation throughout the central nervous system.

*Results:* Consistent with current understanding, wild-type expression of Na_V_1.1 transcript and protein were absent in knock-out mice in the prefrontal cortex, striatum, hippocampus, thalamus, and cerebellum. Increased GFAP was detected in the brain only in the hippocampus. Transcript and protein were detected in wild-type cervical, thoracic and lumbar spinal cord but not in knock-out mice. Unexpectedly, GFAP was increased in all three spinal regions. Therefore, we proceeded to perform transcriptomic analysis of cortex, hippocampus and spinal cord. Pathways associated with monooxygenase activity, fatty acid ligases and lactate transporters were highly dysregulated in the spinal cord.

*Conclusion:* The existence and relevance of pathology of the spinal cord in Dravet syndrome has received scant attention. Our findings are consistent with some systemic symptoms of Dravet syndrome, with the benefits of treatments which may modulate the astrocyte-neuron lactate shuttle such as Stiripentol and ketogenic dietary regimes, and with the efficacy of intrathecal delivery of therapeutics.

**Key Points:**

- Decrease of endogenous *Scn1a* and Na_V_1.1 expression in *Scn1a*^-/-^ mice has a widespread impact on the gene expression profile in the spinal cord.
- Increased GFAP expression observed in the spinal cord of *Scn1a*^-/-^ mice.
- Differentially expressed genes related to monooxygenase activity, fatty acid ligases and lactate transporters in cervical spinal cord of *Scn1a*^-/-^ mice.

## 1. Introduction

Dravet syndrome (DS) is a severe childhood genetic epilepsy, which manifests in the first year or life, accompanied by generalized tonic or clonic seizures, development delay (1, 2), motor dysfunction, ataxia (3) impaired language acquisition (4), sudden unexpected death from epilepsy (SUDEP) and other comorbidities such as hyperactivity and autistic spectrum disorders (5, 6). Approximately 80% of cases are caused by an haploinsufficiency in *SCN1A* which encodes the voltage-gated sodium ion channel, Na_V_1.1 (6).

Rodent models for DS have been characterised extensively to recapitulate the disease, exhibiting phenotypes such as spontaneous seizures, ataxia, hyperactivity, anxiety-like behaviours, low sociability, cognitive deficits and SUDEP (7–10). The knock-out (*Scn1a*^-/-^) Dravet mouse model exhibits spontaneous seizures and ataxia at approximately postnatal day 10 (P10), until death at P14-P16 on 129SvJ and mixed 129SvJ+B6 background (11, 12).

Na_V_1.1 expression is located in the axon initial segment and nodes of Ranvier of the grey matter, and expressed in the hippocampus (CA3 and CA2 regions), dentate gyrus, layer V/VI of the cortex, granular layer, molecular layer, deep nuclei, and Purkinje cells of the cerebellum (13–15). In rodents Na_V_1.1 expression has been detected in the cortex, dentate gyrus, CA1-3 hippocampus and Purkinje cells of the cerebellum (16, 17). Mouse models for DS suggest loss of *Scn1a* is particularly associated with dysfunction of GABAergic interneurons (16) but more recently effects have been seen in CA1 pyramidal cells (18, 19).

The role of Na_V_1.1 outside the brain has been less well studied, but some evidence suggests spinal and peripheral neurons are impacted and this may contribute to the clinical phenotypes in children with DS. *SCN1A* expression has been observed in the cervical, thoracic and lumbar regions of the spinal cord of non-human primates (20). *Scn1a* has also been reported throughout the rodent spinal cord, including the dorsal horn, although at lower levels in the lamina II of the dorsal horn (21) and dorsal root ganglion neurons (21, 22, 22, 23). Deficiency of Na_V_1.1 in the spinal cord has been associated with arthrogryposis multiplex congenita, a disease associated to motor development (24), suggesting this channel may play a role in normal function of the spinal circuit in humans.

While these findings for Na_V_1.1 expression in the spinal cord are intriguing, there is no specific correlation of spinal cord Na_V_1.1 expression to the disease manifestation or symptom onset in DS, specifically ataxia and gait deterioration. To address this, we sought to analyse the spinal cord tissue from the *Scn1a*^-/-^ DS mouse model and specifically assess the *Scn1a* and Na_V_1.1 expression profile, transcriptome assessment profile and spinal cord pathology.

Our findings demonstrate that endogenous *Scn1a* and Na_V_1.1 is significantly decreased in the *Scn1a*^-/-^ spinal cord. We observe a significant increase of GFAP (pan-reactive astrocytic marker; glial fibrillary acidic protein) throughout the spinal cord of *Scn1a*^-/-^ mice and RNA-seq transcriptome analysis revealed significant alterations of gene expression specifically related to monooxygenase pathways, fatty acid ligase and lactate transporters in the cervical spinal cord of *Scn1a*^-/-^ Dravet syndrome mice.

## 2. Materials and Methods

### Animal welfare

All animal studies were conducted under UK Home Office license and were approved by the University College London (UCL) ethical review committee. Animals were housed in individually ventilated cages. 129S-Scn1a^tm1Kea^/Mmjax mice were obtained from the Jackson laboratory (USA) and male mice were crossed with female wild-type CD1 (Charles River, UK). The resulting heterozygous mice were mated to obtain F2 CD1:129Sv Scn1a^-/-^. Weights were measured every other day. Genotyping was performed following the protocol previously described (25).

### Behavioural analysis

All behavioural analysis were blinded and randomised. Righting reflex procedure was performed as previously described (26, 27). The test was video documented and analysed using BORIS (Behavioural Observation Research Interactive Software, http://www.boris.unito.it) v7 (28).

Open field analysis (29) was performed by placing the mice into a grey box (25x25x30cm), motor activity was recorded for 5 minutes and analysed using SMART Video Tracking System v3.0 (Panlab SLU, Spain).

### Tissue Preparation

Mice were euthanized by terminal anaesthesia, followed by cardiac perfusion with Phosphate-buffered saline (PBS). For each organ, the tissues were split in half, for RNA/protein extraction or fixed in 4% paraformaldehyde solution for 48 hours, then transferred to 30% sucrose solution for storage at 4ºC.

### RNA isolation

Total RNA was extracted from the samples using RNeasy mini kit (Qiagen), as per the manufacturer’s instructions. Reverse-transcription was performed using the High-Capacity cDNA Reverse Transcription kit (Applied Biosystems). The resulting cDNA was used for quantitative PCR (qPCR).

### qPCR

Primers and probes were designed targeting the *Scn1a* and *Gapdh* genes. For *Scn1a* forward: TCAGAGGGAAGCACAGTAGAC, reverse: TTCCACGCTGATTTGACAGCA, probe: CCAGAAGAAACCCTTGAGCCCGAA (fluorophore: ABY, quencher: QSY, sourced from ThermoFisher Scientific). For Gapdh forward: ACGGCAAATTCAACGGCAC, reverse: TAGTGGGGTCTCGCTCCTGG, probe: TTGTCATCAACGGGAAGCCCATCA (fluorophore: VIC, quencher: QSY, sourced from ThermoFisher Scientific). 5mL of each cDNA sample was added to a Luna® Universal Probe qPCR Master Mix (NEB). 250nM of primers and probes were added to master mix. The reaction was performed in an QuantStudio 3 (Applied Biosystems). Data was obtained using the software QuantStudio Design and Analysis v1.4 (Applied Biosystems).

### RNA-Seq

Total RNA from the cortex, hippocampus and cervical spinal cord from P14 *Scn1a*^+/+^, *Scn1a*^+/-^ and *Scn1a*^-/-^ was used to for RNA-seq analysis. Samples were processed for using NextSeq 1000/2000 P2 Reagents (Illumina, San Diego, CA). Samples were sequenced on the Illumina HiSeq 2000 platform in the UCL Genomics Facility. Sequence data quality was assessed prior to mapping by using the FASTQC toolkit (http://www.bioinformatics.babraham.ac.uk/projects/fastqc/). Sequenced cDNA libraries were mapped at the transcript and gene levels to the mouse genome (assembly GRCh38) using Salmon v0.8.2. We normalised and compared gene expression using the R package DESeq2 (30), used to test for differential gene expression on log2 normalised gene counts. Analysing clusters for biological enrichment using pathway over-representation analysis was performed using the enrich GO function in the cluster Profiler package for R (31), classifying genes into Gene Ontology (GO) terms for their associated molecular function. Significantly enriched pathways were denoted by FDR corrected p-value <0.05. All statistical tests were two tailed, alpha <0.05 was used to determine statistical significance.

### Free-Floating Immunohistochemistry

40µm sections were cut using a Micron HM 430 freezing microtome (Thermo Fisher Scientific). To analyse CD68 & GFAP immunoreactivity free-Floating Immunohistochemistry was performed following previously described protocol (32, 33). Tissue images were captured using a stereoscopic fluorescence microscope (DM4000; Leica). Quantitative image analysis was performed using Image-Pro Premier v10 (Media Cybernetics).

### Western Blot

Tissues were homogenised in a 2% SDS buffer (10mM Tris HCl pH 7.4, 1mM sodium fluoride, 0.1mM sodium orthovanadate, 2% SDS and 1X complete™, EDTA-free Protease Inhibitor Cocktail) using a tissue lyser (30 Hz for 3 minutes), then lysates were incubated on ice for 20 minutes before centrifuged at 15,000g at 4°C for 15 minutes. Supernatants were kept and the concentrations of resulting protein lysates was determined using the BioRad DC protein assay according to manufacturer’s instructions. Na_V_1.1 western blots used 4-20% Tris-Glycine gels and PageRuler™ Plus Prestained Protein Ladder. Samples were run at 150V for 60 minutes and transferred to 0.45μm PVDF membranes at 70V for 150 minutes. Membranes were blocked in 5% milk in PBS for 1 hour at room temperature. Primary antibodies (Na_V_1.1 Merck, AB5204, 4μg/ml and alpha tubulin Proteintech, 66031-1-Ig, 0.08μg/ml) were diluted in 3% BSA in 0.1% PBST and incubated overnight at 4°. Secondary antibodies (IRDye 800CW goat anti-mouse IgG LiCor, 926-32210, 0.5μg/ml and IRDye 800CW goat anti-rabbit IgG LiCor, 926-32211, 0.5μg/ml) were diluted in the same buffer and incubated for 1 hour at room temperature. Western blots were imaged using the BioRad ChemiDoc MP. Quantification of protein signals was achieved using BioRad Image Lab 6.1 software. The blot images were adjusted by curves in Adobe Photoshop 2024.

### Statistical Analysis

qPCR data of gene expression, western blot protein expression and semi-quantitative immunohistochemical staining analysis were analysed by One-Way ANOVA with Tukey’s multiple comparisons test. Survival data was analysed using the Log-rank (Mantel-Cox) test. Weight and behavioural data were analysed with Mixed-effects model with Tukeys or Dunnett’s multiple comparison test respectively (GraphPad software v9.4.0).

## 3. Results

### Endogenous Na_V_1.1 and *Scn1a* quantification in *Scn1a*^-/-^ spinal cords

We crossed CD1 outbred mice with 129Sv *Scn1a*^-/-^ to generate CD1x129Sv *Scn1a*^-/-^ hybrids, which developed severe ataxia, spontaneous seizures (Supplementary video 1), progressive motor deterioration (Supplementary video 2) and premature death by P14 (Supplementary Figure 1), as previously described in *Scn1a*^-/-^ mouse models (16). At P14 we observed strong Na_V_1.1 protein expression in the hippocampus, cervical, thoracic and lumbar spinal cord regions of wild-type mice; which was reduced in *Scn1a*^+/-^ mice, and not detectable above baseline in *Scn1a*^-/-^ mice (Figure 1A-D; Supplementary Figure 4). RT-qPCR was performed to assess the endogenous *Scn1a* expression and significant difference of *Scn1a* was observed in *Scn1a*^-/-^ hippocampal, lumbar and thoracic spinal cord regions compared to *Scn1a*^+/+^ controls (Figure 1E-H). Endogenous *Scn1a* expression was also decreased throughout the brain and heart of *Scn1a*^-/-^ DS mice (Supplementary Figure 2A-D).

**Figure 1.**
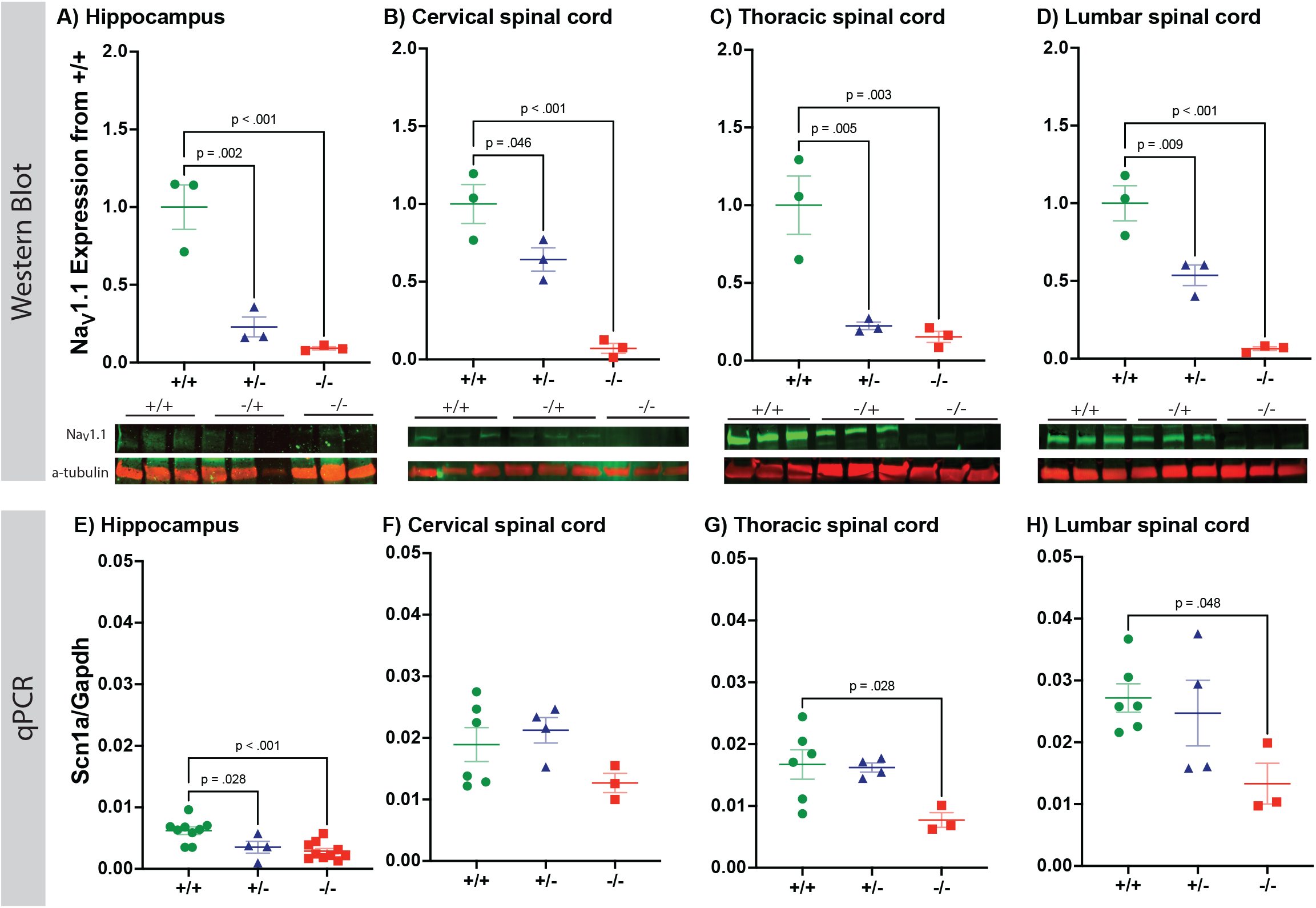
Reduced Na_V_1.1 and *Scn1a* expression in the Spinal Cord of *Scn1a*^-/-^ DS mice. **A)** The hippocampus had a significant decrease in Na _V_1.1 in both *Scn1a* ^+/-^ and *Scn1a* ^-/-^ compared to *Scn1a* ^+/+^ littermates. **B)** The cervical spinal cord had a significant decrease in Na _V_1.1 in both Scn1a^+/-^ and Scn1a^-/-^ compared to Scn1a^+/+^. Similarly, **C)** in the thoracic and **D)** lumbar spinal cord there was a significant decrease in Na_V_1.1 expression in both *Scn1a* ^+/-^ and *Scn1a* ^-/-^ mice. The corresponding quantified blot is at the bottom of each graph and Na 1.1 expression was normalised to Scn1a^+/+^ levels for quantification. n=3 for all groups. One-Way ANOVA with Dunnett’s multiple comparisons test was performed. **E)** The hippocampus revealed significant reduction in endgoenous *Scn1a* expression in Scn1a^-/-^ and *Scn1a* ^+/-^ compared to *Scn1a* ^+/+^. **F)** In the cervical spinal cord, *Scn1a* showed a trend towards decreased expression in *Scn1a* ^-/-^ compared to *Scn1a* ^+/+^. **G)** In the thoracic and **H)** lumbar spinal cord there was a significant decrease in *Scn1a* expression in *Scn1a* ^-/-^ mice compared to *Scn1a* ^+/+^. One-Way ANOVA with Tukey’s multiple comparisons test was performed. n=6 *Scn1a* ^+/+^, 4 *Scn1a* ^+/-^ and 3 *Scn1a* ^-/-^. Graphs are Mean ± SEM.

### Increased GFAP expression in the hippocampus and spinal cord of *Scn1a*^-/-^ mice

Previous studies have demonstrated upregulation of GFAP in the hippocampus of DS mouse models (34, 35). We quantified GFAP expression in samples spanning the CNS from the prefrontal cortex to the lumbar spinal cord. In *Scn1a*^-/-^ mice, GFAP expression was significantly increased only in the hippocampus (Figure 2C) of the brain. Interestingly, GFAP was also increased in cervical, thoracic and lumbar spinal cord (Figure 2F-H).

**Figure 2.**
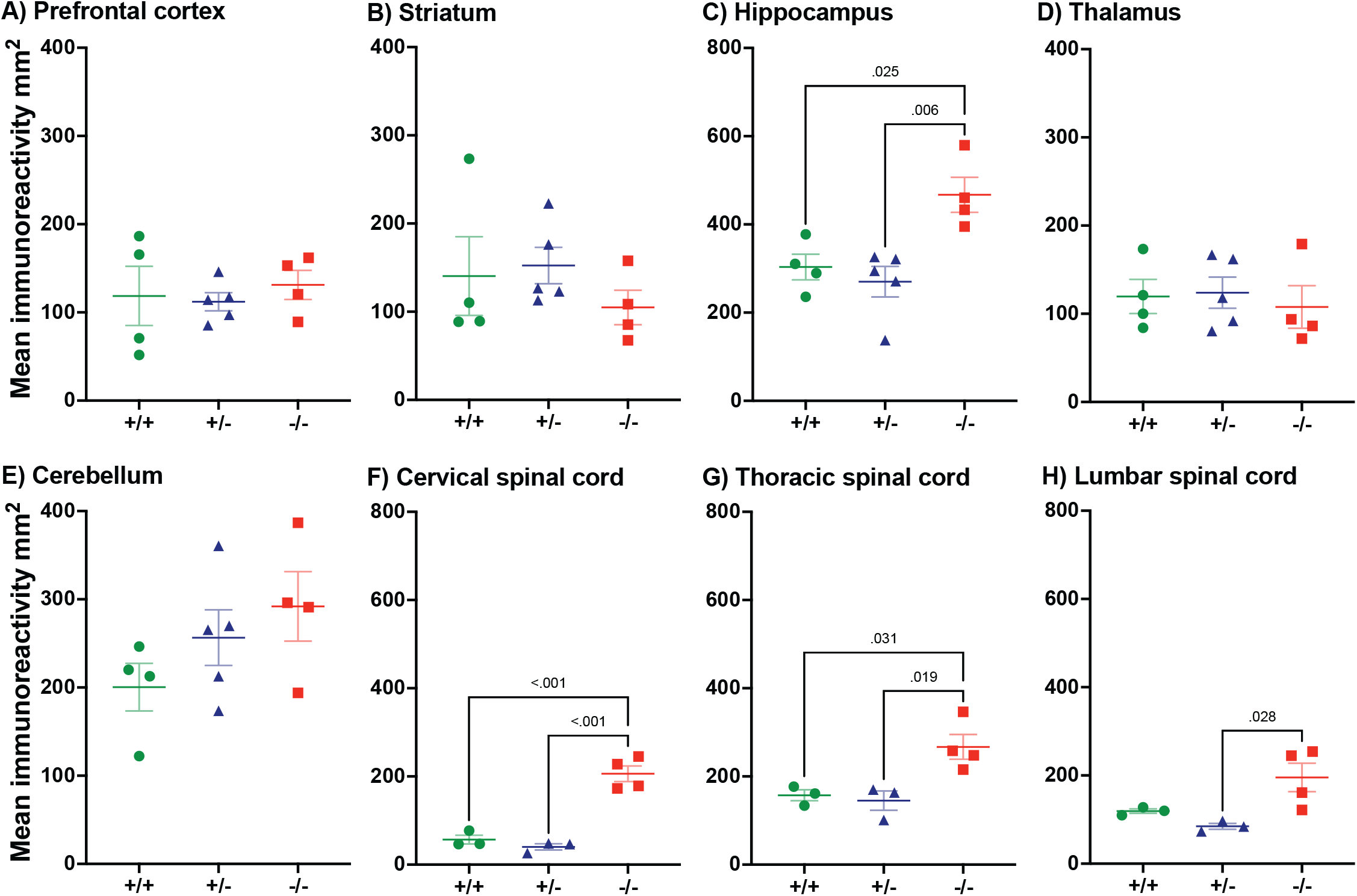
GFAP immunohistochemical analysis in the brain and spinal cord of *Scn1a*^-/-^ DS mice. GFAP immunohistochemical analysis in **C)** the hippocampus showed a significant increase in GFAP in *Scn1a* ^-/-^ mice compared to both *Scn1a* ^+/+^ and *Scn1a* ^+/-^. No statistical difference was observed in the **A)** Prefrontal cortex, **B)** striatum, **D)** thalamus and **E)** cerebellum. **F)** The cervical spinal cord, which showed a significant increase in GFAP in *Scn1a*^-/-^ compared to both *Scn1a* ^+/+^ and *Scn1a* ^+/-^. **G)** Similar results were found in the thoracic region. **H)** In the lumbar spinal cord, we observed a signficant increase in GFAP in *Scn1a* ^-/-^ compared to *Scn1a* ^+/-^. n=3/5 for all groups. One-Way ANOVA with Tukey’s multiple comparisons test was performed. Graphs are Mean ± SEM.

### Altered gene expression in the cortex, hippocampus and cervical spinal cord of *Scn1a*^-/-^ mice

To provide an unbiased global assessment of gene expression, we performed RNA-seq on isolated cortex, hippocampus and cervical spinal cord tissue (P14 mice, n=3/4 for each group). This revealed a wide array of differentially expressed genes (DEGs) in *Scn1a*^-/-^ compared to *Scn1a*^+/+^ in all tissues.

In the hippocampus, there were 311 down-regulated DEGs and 538 up-regulated DEGs in *Scn1a*^-/-^ compared to *Scn1a*^+/+^. GO categories for DEGs of the hippocampus were associated with cell proliferation, cytoskeletal regulation, neurotransmitter regulation, voltage gated sodium channels, transcription, growth factor activity, extracellular matrix assembly and glycosylation (Figure 3A-C). Similar DEGs were noted for the cortex (Figure 5A-C).

**Figure 3.**
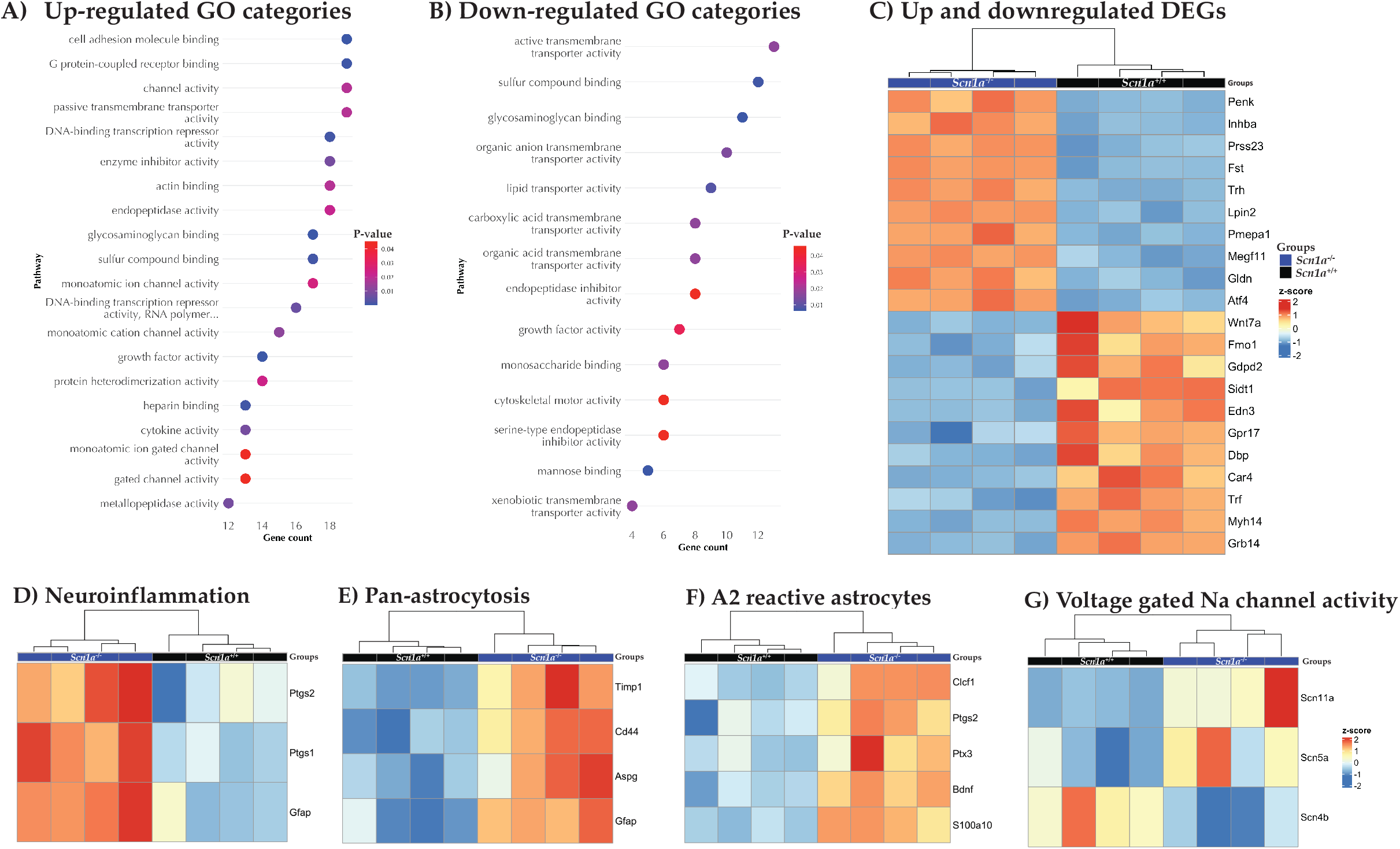
Altered gene expression in the *Scn1a*^-/-^ hippocampus. **A)** Up-regulated and **B)** down-regulated pathways in Scn1a^-/-^ mice compared to *Scn1a*^+/+.^ **C)** Heatmap showing the top 10, up and down-regulated DEGs in *Scn1a*^-/-^ and *Scn1a* ^+/+^ hippocampus. Heatmaps showing DEGs for markers of **D)** neuroinflammation, **E)** pan-astrocytosis, **F)** A2 reactive astrocytes and **G)** voltage-gated Na channel activity between Scn1a^-/-^ and Scn1a^+/+^ in the hippocampus. n=4 for each group.

We observed an up-regulation of pan-astrocytic markers in the *Scn1a*^-/-^ cortex and hippocampus and DEG associated with “A2” reactive astrocytes which promote pro anti-inflammatory factors to protect neurons (Figure 3F & Figure 5G). Conversely, in the cortex, we noted DEG associated with “A1” reactive astrocytes, which activate pro-inflammatory substances for neuronal death (Figure 5F). We also noted DEG associated with voltage-gated sodium channels in the hippocampus (Figure 3G) and cortex (Figure 5H).

In the cervical spinal cord, there were 65 DEGs (22 up-regulated and 43 down-regulated, Figure 4B) in *Scn1a*^-/-^ compared to *Scn1a*^+/+^, the GO categories for up-regulated genes were associated with oxidation-reduction pathways, monooxygenase activity, lactate transporters and iron metabolism (Figure 4A). However, in contrast to our immunohistochemical data showing upregulation of GFAP expression, we did not observe GO categories associated with activation of astrocytes. Figure 6 shows the overlap of down & up-regulated GO categories and DEGs in the cortex, hippocampus, and cervical spinal cord.

**Figure 4.**
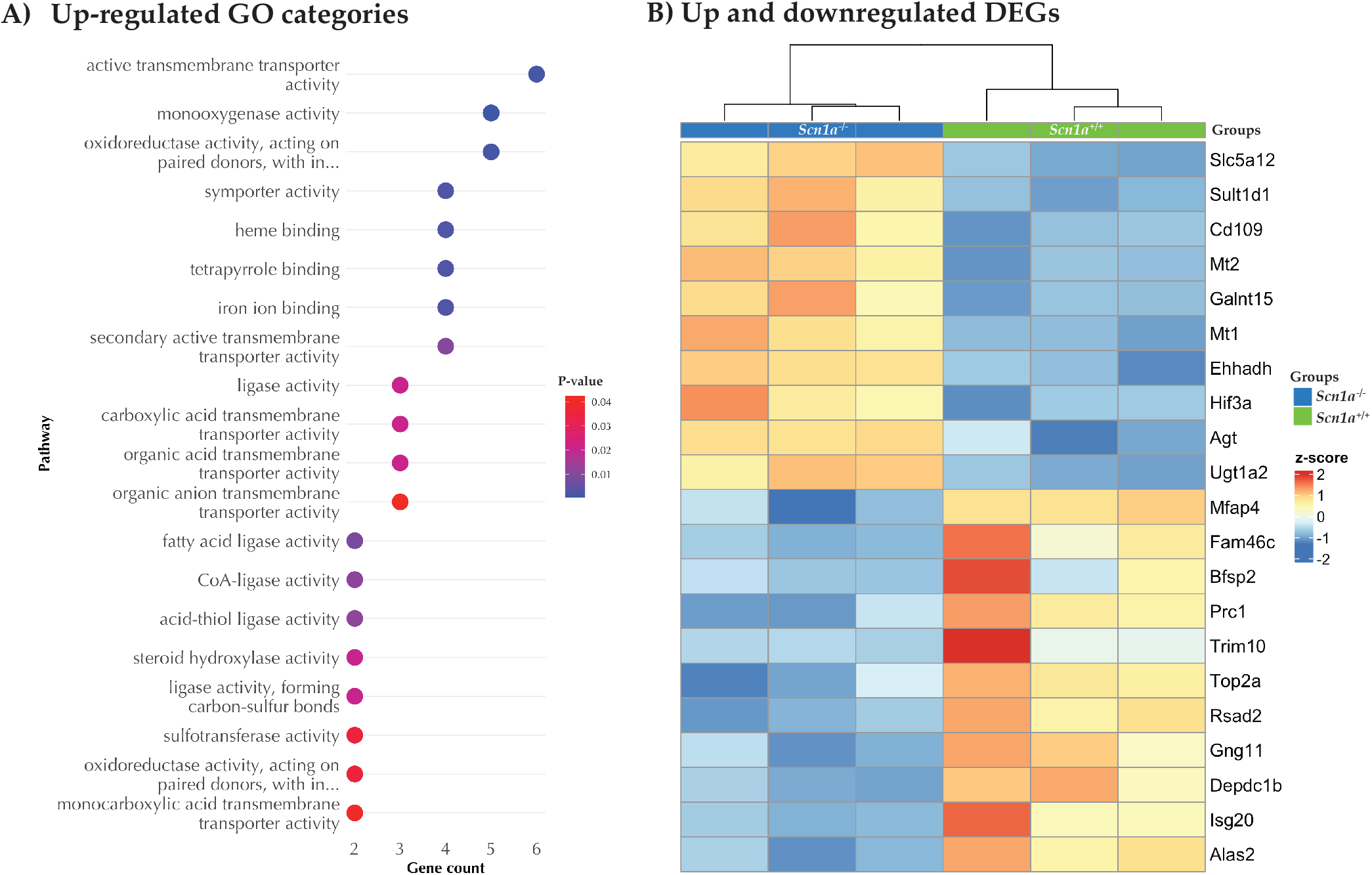
Altered gene expression in *Scn1a*^-/-^cervical spinal cord. **A)** Pathways associated with up-regulated genes in the *Scn1a* ^-/-^ compared to *Scn1a* ^+/+^ cervical spinal cord. **B)** Heatmap showing the top 10 up-regulated and down-regulated DEG’s in Scn1a^-/-^ cervical spinal cord. n=3 for each group. For two categories (top to bottom) their full text is; “oxidoreductase activity, acting on paired donors, with incorporation or reduction of molecular oxygen” and “oxidoreductase activity, acting on paired donors, with incorporation or reduction of molecular oxygen, NAD(P)H as one donor, and incorporation of one atom of oxygen”.

**Figure 5.**
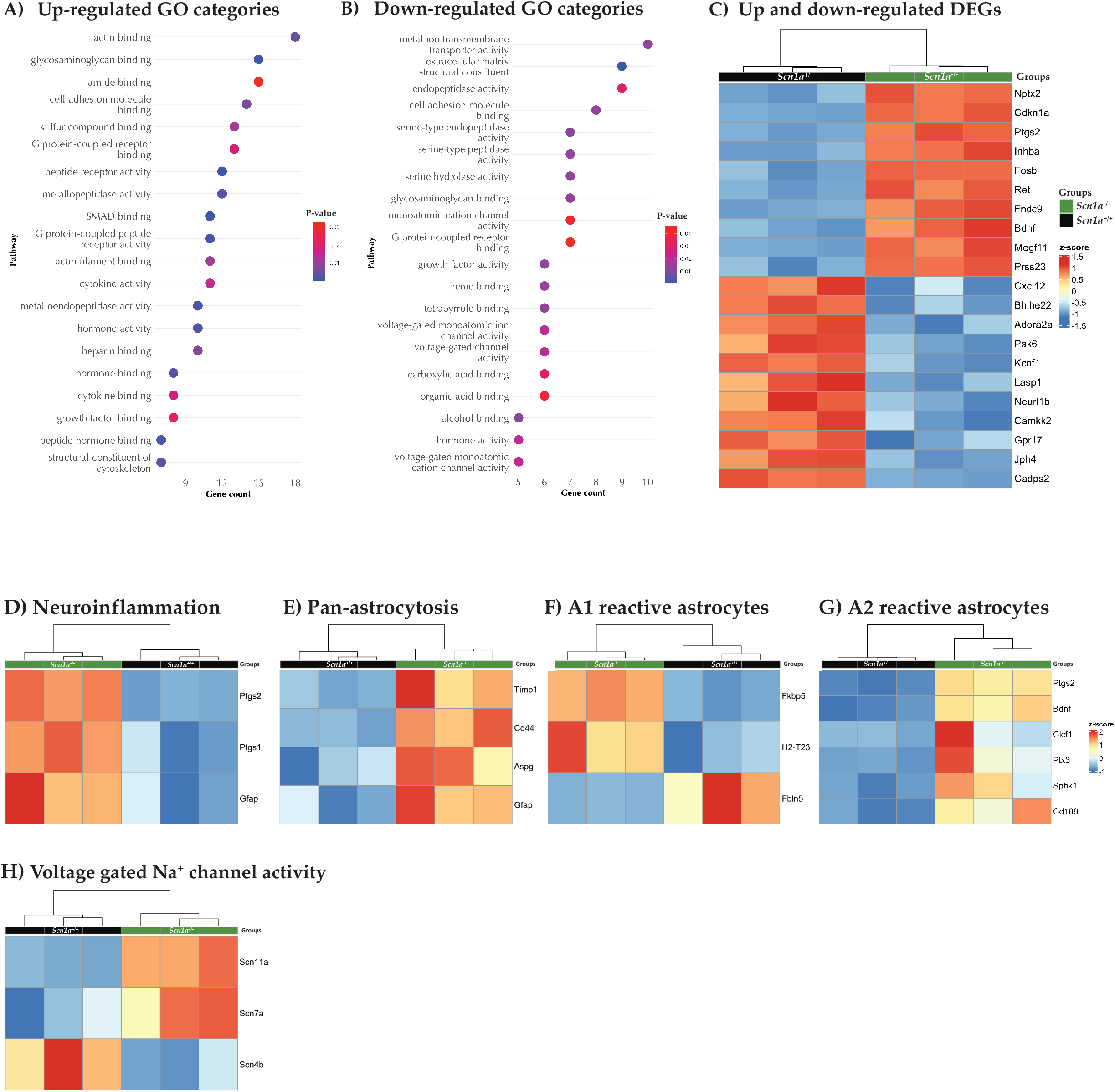
*A*ltered gene expression in *Scn1a*^-/-^ cortex. **A)** Up-regulated and **B)** down-regulated pathways in the *Scn1a* ^-/-^ compared to *Scn1a* ^+/+^. **C)** Heatmap showing the top 10 DEGs in *Scn1a* ^-/-^ and *Scn1a* ^+/+^ cortex. Heatmaps showing DEG for markers of **D)** neuroinflammation, **E)** pan-astrocytosis, **F)** A1 reactive astrocytes, **G)** A2 reactive astrocytes and **H)** voltage-gated Na^+^ channel activity between *Scn1a* ^-/-^ and *Scn1a* ^+/+^ cortex. n=3 for each group.

**Figure 6.**
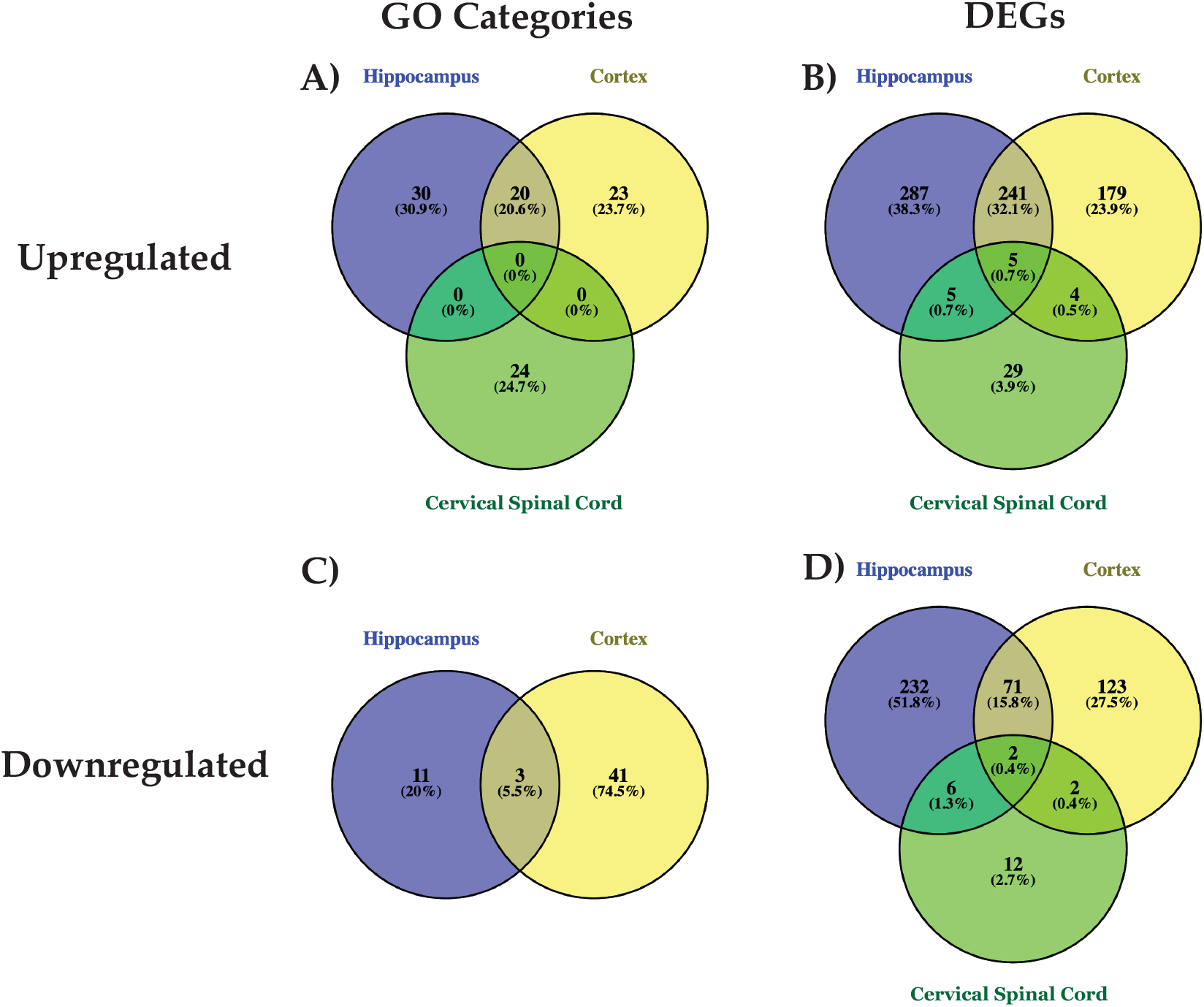
Number of shared GO terms and DEGs between the studies regions. The number of up and down-regulated GO terms (**A** and **C**) and differentially expressed genes (DEGs, **B** and **D**) for each region analysed in RNAseq. The hippocampus and cortex share a number of GO categories and DEGs, while the spinal cord has no common GO categories with the other two brain regions. Among the DEGs, less than 2% of all the significantly up or downregulated genes were in the cervical spinal cord overlapped with cortex and hippocampus.

## 4. Discussion

In this study we have characterised extensive changes in *Scn1a* and Na_V_1.1 expression throughout the spinal cord of *Scn1a*^-/-^ DS mice, which confirms and extends previous studies (36, 37). We have also demonstrated significant differences in *Scn1a* expression in the heart, brain of *Scn1a*^-/-^ mice, and most importantly in the spinal cord (38).

We observed a significant increase of GFAP in the hippocampus of *Scn1a*^-/-^ DS mice, which has previously seen in the CNS of DS mouse models (34, 35). More importantly we have shown for the first time that GFAP expression is increased within the spinal cord of *Scn1a*^-/-^ DS mice. Interestingly, changes in GFAP expression in the spinal cord correlated with the decrease in Na_V_1.1 expression. Other studies have shown that restoring Na_V_1.1 expression in the brain reduced inflammatory markers (39), suggesting a two-way relationship between the levels of Na_V_1.1 and GFAP, that may be conserved across the CNS. These findings are important as mechanism of many of the motor phenotypes associated with DS are not clear, and our data along with other studies, suggests reactive astrocytes may contribute to ataxia, gait disturbances (40), cardiac autonomic function dysautonomia that characterise DS (41).

Altered gene expression profiles have been reported in epilepsies and in models of Dravet syndrome, specifically in the brain (25, 39, 42). For the first time we have conducted gene expression analysis in the cervical spinal cord in *Scn1a*^-/-^ DS mouse model. Interestingly the cervical spinal cord analysis demonstrated distinctive dysregulation of GO categories and DEGs in comparison to the hippocampus and the cortex. We only noted approximately 1.3% of up or down regulated DEG’s in the cervical spinal cord overlapping with the hippocampus and cortex. In contrast the hippocampus and cortex shared a number of GO categories and DEG’s.

In the hippocampus, our transcriptome analysis was consistent with previous studies conducted on *Scn1a*^+/-^ DS mouse models which also highlighted GO categories of DEG relevant to DS phenotypes, including cytoskeletal regulation, extracellular matrix assembly and voltage gated sodium channels (39). Markers for astrogliosis and A2 reactive astrocytes were also up-regulated in the *Scn1a*^-/-^ hippocampus (34, 39, 43). For the first time we report up-regulation of pan-astrocytic, A1 and A2 reactive astrocyte markers in the cortex of *Scn1a*^-/-^ DS mice, correlating with previous studies conducted on *Scn1a*^+/-^ DS mouse models (39). However, DEG’s associated with neuroinflammation or reactive astrocytes were not detected in the cervical spinal cord.

In the *Scn1a*^+/-^ cervical spinal cord we observed changes in the expression of genes regulating metabolic homeostasis, including those associated to the family of cytochrome P450 enzymes (CYP). Changes in metabolic homeostasis play a major role in neuronal excitability and seizure susceptibility and could provide a new insight into pathogenesis as well as current and potential treatments for DS. CYP are monooxygenases and have been reported to metabolise anti-seizure medications for DS such as phenobarbital and topiramate and thus CYP reduces the anti-seizure medications half-life and action (44). These findings suggest a possible mechanism for the drug resistance observed in DS patients (45).

Genes associated with fatty acid ligase and CoA ligase activity were also up-regulated in *Scn1a*^-/-^ cervical spinal cord. These pathways are involved in producing ketone bodies in astrocytes from ketogenic diets, resulting in production of ATP in the mitochondria, increasing energy reserves and providing neuroprotection (46). This may have a critical role in understanding the therapeutic effect of ketogenic diets in DS patients. Interestingly, lactate transporter gene, *Slc5a12* was also up-regulated, this may modulate the astrocyte-neuron lactate shuttle, where lactate formed through lactate dehydrogenase by the astrocytes is used to support energy demands by hyperactive neuronal networks. This is consistent with previous suggestions that seizures induce lactate production (47).

All these pathways also give new insight into the mechanism of action of Stiripentol, anti-seizure medication for in DS. Stiripentol has been shown to inhibit the function of many CYPs and is thought to act by inhibiting lactate dehydrogenase (48) and more importantly previous studies have indicated ataxia as an adverse event in DS patients after Stiripentol treatment (49). Thus, this may regulate astrocyte-neuron lactate shuttle, which we have now shown is disrupted in *Scn1a*^-/-^ cervical spinal cord.

In conclusion, our findings provide new potential mechanisms for ataxia, gait disturbances and dysautonomia present in DS patients. In addition, we observed DEG associated with CYP, fatty acid ligase, CoA ligase and lactate transporters in the cervical spinal cord. Our data provide new insight into the pivotal role of astrocyte-neuron lactate shuttle modulated by Stiripentol and ketogenic dietary regimes. More importantly these findings highlight the possible mechanisms influencing efficacy of intrathecal delivery of therapies for DS such as Stoke therapeutics, Phase 1 STK-001 treatment (50).

## Acknowledgements

RK, SNW, SS received funding from MRC DPFS grant MR/P026494/1. RK, SNW, SS, JAD received funding from LifeArc Philanthropic fund, P2020-0008. RK, SNW, SS, EC received funding from Great Ormond Street Hospital Charity grant, V4720. JAD received funding from CONICYT Becas Chile Doctoral Fellowship program 72160294.

RK, SNW and JAD developed experimental design. JAD, EC, AK, SS, SNW & RK contributed to data. JAD, SS, SNW & RK contributed to manuscript.

## References

1. Dravet C. The core Dravet syndrome phenotype. Epilepsia. 2011;52:3–9.

2. Scheffer IE, Berkovic S, Capovilla G et al. ILAE classification of the epilepsies: Position paper of the ILAE Commission for Classification and Terminology. Epilepsia. 2017;58:512–521.

3. Catarino CB, Liu JYW, Liagkouras I et al. Dravet syndrome as epileptic encephalopathy: evidence from long-term course and neuropathology. Brain. 2011;134:2982–3010.

4. Genton P, Velizarova R, Dravet C. Dravet syndrome: the long-term outcome. Epilepsia. 2011;52 Suppl 2:44–49.

5. Ragona F, Granata T, Dalla Bernardina B et al. Cognitive development in Dravet syndrome: a retrospective, multicenter study of 26 patients. Epilepsia. 2011;52:386–392.

6. Brunklaus A, Ellis R, Reavey E, Forbes GH, Zuberi SM. Prognostic, clinical and demographic features in SCN1A mutation-positive Dravet syndrome. Brain. 2012;135:2329–2336.

7. Han S, Tai C, Westenbroek RE et al. Autistic-like behaviour in Scn1a+/-mice and rescue by enhanced GABA-mediated neurotransmission. Nature. 2012;489:385–390.

8. Ito S, Ogiwara I, Yamada K et al. Mouse with Nav1.1 haploinsufficiency, a model for Dravet syndrome, exhibits lowered sociability and learning impairment. Neurobiology of Disease. 2013;49:29–40.

9. Kalume F, Westenbroek RE, Cheah CS et al. Sudden unexpected death in a mouse model of Dravet syndrome. Journal of Clinical Investigation. 2013;123:1798–1808.

10. Kearney J. Sudden unexpected death in dravet syndrome. Epilepsy currents. 2013;13:264–265.

11. Kalume F, Yu FH, Westenbroek RE, Scheuer T, Catterall WA. Reduced sodium current in Purkinje neurons from Nav1.1 mutant mice: implications for ataxia in severe myoclonic epilepsy in infancy. J Neurosci. 2007;27:11065–11074.

12. Patra PH, Serafeimidou-Pouliou E, Bazelot M, Whalley BJ, Williams CM, McNeish AJ. Cannabidiol improves survival and behavioural co-morbidities of Dravet syndrome in mice. Br J Pharmacol. 2020

13. Whitaker WR, Clare JJ, Emson PC. Differential distribution of voltage-gated sodium channel alpha-and beta-subunits in human brain. Annals of the New York Academy of Sciences. 1999;868:88–92.

14. Whitaker WR, Clare JJ, Powell AJ, Chen YH, Faull RL, Emson PC. Distribution of voltage-gated sodium channel alpha-subunit and beta-subunit mRNAs in human hippocampal formation, cortex, and cerebellum. The Journal of comparative neurology. 2000;422:123–139.

15. Whitaker WR, Faull RL, Waldvogel HJ, Plumpton CJ, Emson PC, Clare JJ. Comparative distribution of voltage-gated sodium channel proteins in human brain. Brain research Molecular brain research. 2001;88:37–53.

16. Yu FH, Mantegazza M, Westenbroek RE et al. Reduced sodium current in GABAergic interneurons in a mouse model of severe myoclonic epilepsy in infancy. 2006;9:1142–1149.

17. Ogiwara I, Iwasato T, Miyamoto H et al. Nav1.1 haploinsufficiency in excitatory neurons ameliorates seizure-associated sudden death in a mouse model of Dravet syndrome. Human Molecular Genetics. 2013;22:4784–4804.

18. Jones SP, O’Neill N, Muggeo S, Colasante G, Kullmann DM, Lignani G. Developmental instability of CA1 pyramidal cells in Dravet Syndrome. bioRxiv. 2022

19. Almog Y, Fadila S, Brusel M, Mavashov A, Anderson K, Rubinstein M. Developmental alterations in firing properties of hippocampal CA1 inhibitory and excitatory neurons in a mouse model of Dravet syndrome. Neurobiol Dis. 2020;148:105209.

20. Hsiao J, Yuan TY, Tsai MS et al. Upregulation of Haploinsufficient Gene Expression in the Brain by Targeting a Long Non-coding RNA Improves Seizure Phenotype in a Model of Dravet Syndrome. EBioMedicine. 2016;9:257–277.

21. Blankenship ML, Coyle DE, Baccei ML. Transcriptional expression of voltage-gated Na+ and voltage-independent K+ channels in the developing rat superficial dorsal horn. Neuroscience. 2013;231:305–314.

22. Ho C, O’Leary ME. Single-cell analysis of sodium channel expression in dorsal root ganglion neurons. Mol Cell Neurosci. 2011;46:159–166.

23. Hildebrand ME, Mezeyova J, Smith PL, Salter MW, Tringham E, Snutch TP. Identification of sodium channel isoforms that mediate action potential firing in lamina I/II spinal cord neurons. Mol Pain. 2011;7:67.

24. Jaber D, Gitiaux C, Blesson S et al. De novo mutations of SCN1A are responsible for arthrogryposis broadening the SCN1A-related phenotypes. J Med Genet. 2021;58:737–742.

25. Miller AR, Hawkins NA, McCollom CE, Kearney JA. Mapping genetic modifiers of survival in a mouse model of Dravet syndrome. Genes, Brain and behavior. 2014;13:163–172.

26. Jänicke B, Coper H. Tests in Rodents for Assessing Sensorimotor Performance During Aging. Advances in Psychology 114. Elsevier; 1996. p. 201–233.

27. Butchbach ME, Edwards JD, Burghes AH. Abnormal motor phenotype in the SMNDelta7 mouse model of spinal muscular atrophy. Neurobiol Dis. 2007;27:207–219.

28. Friard O, Gamba M. BORIS: a free, versatile open-source event-logging software for video/audio coding and live observations. Methods in Ecology and Evolution. 2016;7:1325–1330.

29. Seibenhener ML, Wooten MC. Use of the Open Field Maze to measure locomotor and anxiety-like behavior in mice. J Vis Exp. 2015 e52434.

30. Love MI, Huber W, Anders S. Moderated estimation of fold change and dispersion for RNA-seq data with DESeq2. Genome Biol. 2014;15:550.

31. Wu T, Hu E, Xu S et al. clusterProfiler 4.0: A universal enrichment tool for interpreting omics data. Innovation (Camb). 2021;2:100141.

32. Karda R, Perocheau DP, Suff N et al. Continual conscious bioluminescent imaging in freely moving somatotransgenic mice. Sci Rep. 2017;7:6374.

33. Karda R, Rahim AA, Wong AMS et al. Generation of light-producing somatic-transgenic mice using adeno-associated virus vectors. Sci Rep. 2020;10:2121.

34. Martín-Suárez S, Abiega O, Ricobaraza A, Hernandez-Alcoceba R, Encinas JM. Alterations of the Hippocampal Neurogenic Niche in a Mouse Model of Dravet Syndrome. Front Cell Dev Biol. 2020;8:654.

35. Satta V, Alonso C, Díez P et al. Neuropathological Characterization of a Dravet Syndrome Knock-In Mouse Model Useful for Investigating Cannabinoid Treatments. Frontiers in Molecular Neuroscience. 2021;13

36. Beckh S, Noda M, Lübbert H, Numa S. Differential regulation of three sodium channel messenger RNAs in the rat central nervous system during development. The EMBO journal. 1989

37. Duflocq A, Le Bras B, Bullier E, Couraud F, Davenne M. Nav1.1 is predominantly expressed in nodes of Ranvier and axon initial segments. Molecular and cellular neurosciences. 2008;39:180–192.

38. Malhotra JD, Chen C, Rivolta I et al. Characterization of sodium channel α-and β-subunits in rat and mouse cardiac myocytes. Circulation. 2001;103:1303–1310.

39. Valassina N, Brusco S, Salamone A et al. Scn1a gene reactivation after symptom onset rescues pathological phenotypes in a mouse model of Dravet syndrome. Nat Commun. 2022;13:161.

40. Rodda JM, Scheffer IE, McMahon JM, Berkovic SF, Graham HK. Progressive gait deterioration in adolescents with Dravet syndrome. Arch Neurol. 2012;69:873–878.

41. Sahai N, Bard AM, Devinsky O, Kalume F. Disordered autonomic function during exposure to moderate heat or exercise in a mouse model of Dravet syndrome. Neurobiology of Disease. 2021;147:105154.

42. Hawkins NA, Calhoun JD, Huffman AM, Kearney JA. Gene expression profiling in a mouse model of Dravet syndrome. Exp Neurol. 2018

43. Hawkins NA, Zachwieja NJ, Miller AR, Anderson LL, Kearney JA. Fine Mapping of a Dravet Syndrome Modifier Locus on Mouse Chromosome 5 and Candidate Gene Analysis by RNA-Seq. PLOS Genetics. 2016;12:e1006398.

44. Brodie MJ, Mintzer S, Pack AM, Gidal BE, Vecht CJ, Schmidt D. Enzyme induction with antiepileptic drugs: cause for concern. Epilepsia. 2013;54:11–27.

45. Sullivan J, Wirrell EC. Dravet Syndrome as an Example of Precision Medicine in Epilepsy. Epilepsy Curr. 2023;23:4–7.

46. Veyrat-Durebex C, Reynier P, Procaccio V et al. How Can a Ketogenic Diet Improve Motor Function. Front Mol Neurosci. 2018;11:15.

47. Purnell BS, Alves M, Boison D. Astrocyte-neuron circuits in epilepsy. Neurobiol Dis. 2023;179:106058.

48. Sada N, Lee S, Katsu T, Otsuki T, Inoue T. Epilepsy treatment. Targeting LDH enzymes with a stiripentol analog to treat epilepsy. Science. 2015;347:1362–1367.

49. Eschbach K, Knupp KG. Stiripentol for the treatment of seizures in Dravet syndrome. Expert Rev Clin Pharmacol. 2019;12:379–388.

50. Therapeutics S. Stoke Therapeutics Announces Positive New Safety & Efficacy Data from Patients Treated with STK-001 in the Phase 1/2a Studies (MONARCH & ADMIRAL) and the SWALLOWTAIL Open-Label Extension (OLE) Study in Children and Adolescents with Dravet Syndrome. 2023. Available from: https://investor.stoketherapeutics.com/news-releases/news-release-details/stoke-therapeutics-announces-positive-new-safety-efficacy-data

