## Supplementary Figures for "Spinal cord pathology in a Dravet Syndrome mouse model"

### Supplementary Figure 1

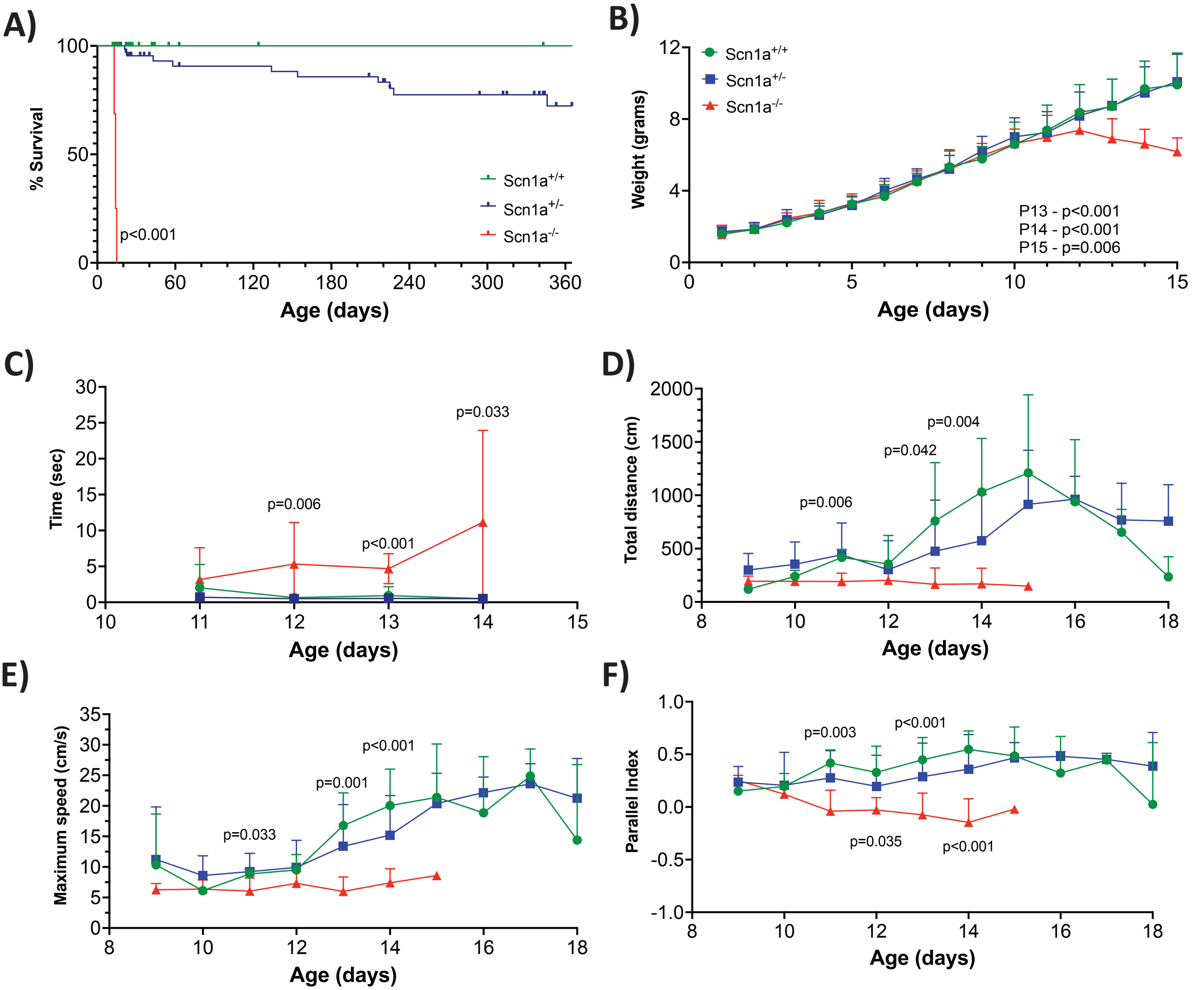


**S. Figure 1. Characterization of the 129Sv x CD1 *Scn1a*$^{-/-}$ DS mouse model. A)** A significant reduction in survival was observed for the *Scn1a* $^{-/-}$ mice compared to *Scn1a* $^{+/+}$ and *Scn1a* $^{+/-}$. **B)** Weight of the *Scn1a* $^{-/-}$ mice was significantly lower than *Scn1a* $^{+/+}$ and *Scn1a* $^{+/-}$ from day 13 , 14 and 15. n; *Scn1a* $^{+/+}$=28, *Scn1a* $^{+/-}$=74 and *Scn1a* $^{-/-}$= 16. **C)** Righting reflex was performed between P11 and P14. *Scn1a* $^{-/-}$ mice displayed significant increase in time to recover from supine to prone position at P12 , P13 and P14 . n; *Scn1a* $^{+/+}$= 36, *Scn1a* $^{+/-}$=22 and *Scn1a* $^{-/-}$= 14. **D)** The total distance walked by *Scn1a* $^{-/-}$ mice was significantly different at P11, 13 and 14 compared to *Scn1a* $^{+/+}$ and *Scn1a* $^{+/-}$. **E)** Maximum speed significantly declined in the *Scn1a* $^{-/-}$ mice at P11, 13 and 14 compared to *Scn1a* $^{+/+}$ littermates. **F)** *Scn1a* $^{-/-}$ mice showed significant changes in direction from P11-14, as seen by the decreasing parallel index; when compared to *Scn1a* $^{+/+}$. n; *Scn1a* $^{+/+}$=8, *Scn1a* $^{+/-}$=28 and *Scn1a* $^{-/-}$= 7. All points are mean ± SD. One-Way ANOVA with Tukey's multiple comparisons test was performed.

### Supplementary Figure 2

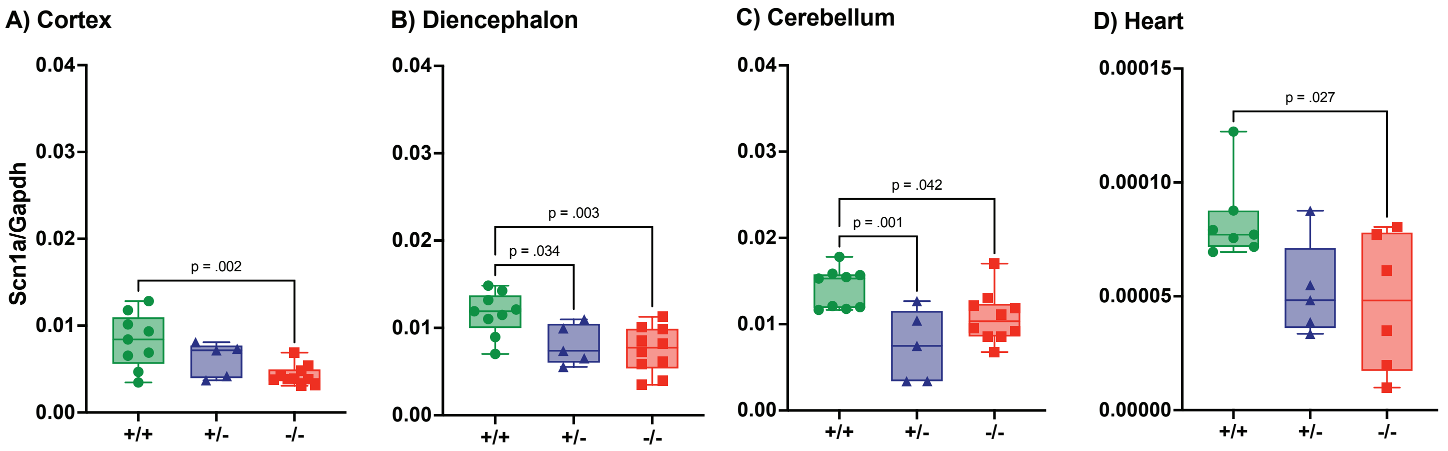


**S. Figure 2. *Scn1a* expression in the brain and heart of *Scn1a*$^{-/-}$ DS mice. A)** In the cortex, endogenous *Scn1a* expression was significantly reduced in the *Scn1a* $^{-/-}$ compared to *Scn1a* $^{+/+}$ mice. **B)** The diencephalic region and **C)** cerebellum showed significant down-regulation in *Scn1a* expression in both the *Scn1a* $^{-/-}$ and *Scn1a* $^{-/+}$ compared to *Scn1a* $^{+/+}$ control group. **D)** The heart showed a significant decrease in *Scn1a* in *Scn1a* $^{-/-}$ compared to *Scn1a* $^{+/+}$. One-Way ANOVA with Tukey's multiple comparisons test was performed. For brian regions; n= 9 *Scn1a* $^{+/+}$, 5 *Scn1a* $^{+/-}$ and 10 *Scn1a* $^{-/-}$. The heart; n= 7 *Scn1a* $^{+/+}$, 5 *Scn1a* $^{+/-}$ and 6 *Scn1a* $^{-/-}$.

### Supplementary figure 3

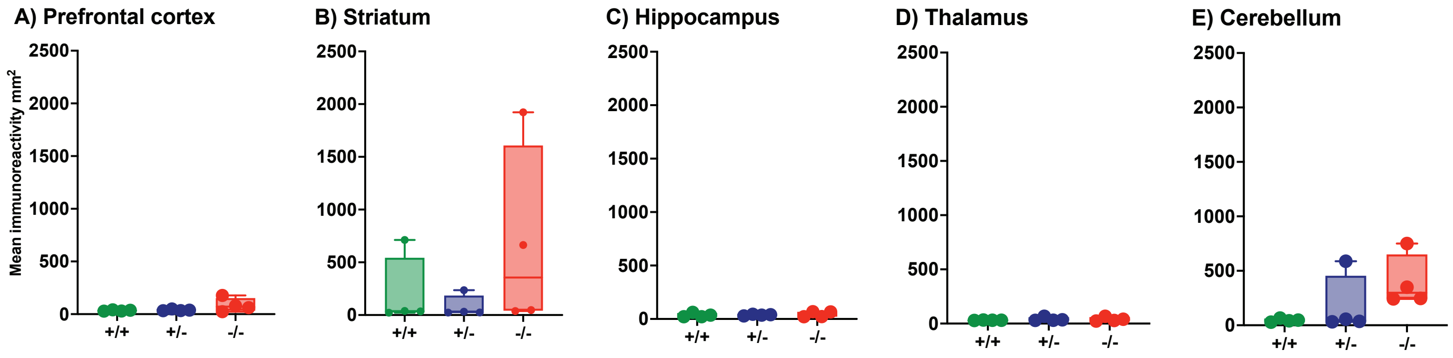


**S. Figure 4. Microglia Immunohistochemical analysis in the brain *Scn1a*$^{-/-}$ DS mice.** CD68 immunohistochemical analysis to detect microglia activity showed no significant difference between the genotypes in each of the discrete brain regions analysed. n=4 for all groups. One-Way ANOVA with Dunnett's multiple comparisons test was performed.

### Supplementary Figure 4

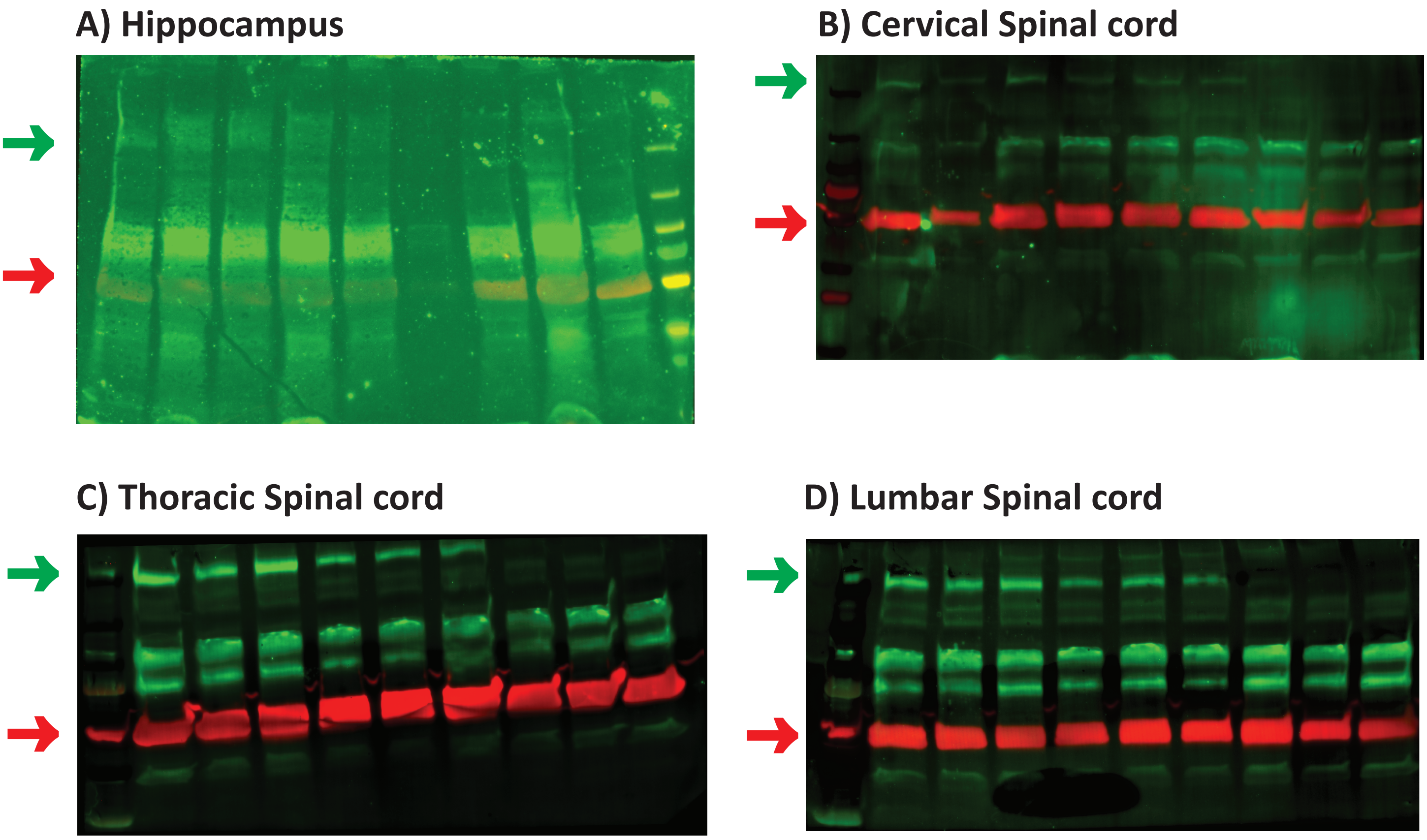


**S. Figure 4. Original Na$_V$1.1 western blot images from hippocampus and spinal cord of *Scn1a*$^{+/+}$, *Scn1a*$^{+/-}$ and Scn1a$^{-/-}$ DS mice.** Arrows show quantified bands; green for Na$_V$1.1 and red for a-tubulin. A) Hippocampus, B) Cervical spinal cord, C) Thoracic spinal cord and D) Lumbar spinal cord. Quantified using BioRad Image Lab software v6.1.0, build 7.
